# Return of intracranial beta oscillations and traveling waves with recovery from traumatic brain injury

**DOI:** 10.1101/2024.07.19.604293

**Authors:** Alex Vaz, Connor Wathen, Stephen Miranda, Rachel Thomas, Timothy Darlington, Rashad Jabarkheel, Samuel Tomlinson, John Arena, Kamila Bond, Sanjana Salwi, Sonia Ajmera, Ludovica Bachschmid-Romano, James Gugger, Danielle Sandsmark, Ramon Diaz-Arrastia, James Schuster, Ashwin G. Ramayya, Iahn Cajigas, Bijan Pesaran, H. Isaac Chen, Dmitriy Petrov

## Abstract

Traumatic brain injury (TBI) remains a pervasive clinical problem associated with significant morbidity and mortality. However, TBI remains clinically and biophysically ill-defined, and prognosis remains difficult even with the standardization of clinical guidelines and advent of multimodality monitoring. Here we leverage a unique data set from TBI patients implanted with either intracranial strip electrodes during craniotomy or quad-lumen intracranial bolts with depth electrodes as part of routine clinical practice. By extracting spectral profiles of this data, we found that the presence of narrow-band oscillatory activity in the beta band (12-30 Hz) closely corresponds with the neurological exam as quantified with the standard Glasgow Coma Scale (GCS). Further, beta oscillations were distributed over the cortical surface as traveling waves, and the evolution of these waves corresponded to recovery from coma, consistent with the putative role of waves in perception and cognitive activity. We consequently propose that beta oscillations and traveling waves are potential biomarkers of recovery from TBI. In a broader sense, our findings suggest that emergence from coma results from recovery of thalamo-cortical interactions that coordinate cortical beta rhythms.

## Introduction

Traumatic brain injury (TBI) is a pervasive clinical phenomenon in the US population, affecting approximately 10% of all individuals (Frost et al. 2013) with much higher estimates in the young adult population (McKinlay et al. 2008). TBI therefore remains an important area of research for improvement of clinical strategies, particularly in the assessment of severity of disease. However, neuroprognostication after TBI remains difficult, complicating efforts towards prognosis, end of life decision making, and the application of neuroprotective therapies. An increased focus towards tractable biomarkers of TBI is therefore underway with several lines of research showing some promise in coagulation and inflammation pathways in both CSF and peripheral blood (Stein et al. 2011; Sharma and Laskowitz 2012; Gan et al. 2019; Diaz-Arrastia et al. 2014). Functional and electrophysiological biomarkers offer a parallel way forward, but it remains unclear which electroencephalographic features are most informative of TBI and its progression (Edlow et al. 2021, 2017; Schiff, Nauvel, and Victor 2014; Laureys and Schiff 2012; Owen, Schiff, and Laureys 2009).

Possible candidates of spectral features related to decreased neurological function broadly focus on the disruption of normal oscillatory electrophysiological activity. For example, intracranial EEG recordings during the induction and emergence from anesthesia demonstrate significant changes in alpha and beta band connectivity across the brain (Weiner et al. 2023; Malekmohammadi et al. 2019). Previous literature using quantitative scalp EEG shows that beta oscillatory power is reduced after cerebral infarcts (Wu et al. 2016; Foreman and Claassen 2012) and in traumatic brain brain injury (Mofakham et al. 2022). Altogether, it may be the case that recovery from coma in traumatic brain injury may be accompanied by a return of physiological beta oscillations known to occur in the human brain. However it largely remains uncharacterized how these dynamics may appear with invasive recordings, and if they may be used to prognosticate recovery and ultimate outcomes in TBI patients.

We therefore hypothesized that the presence and recovery of intracranial beta oscillations may be tractable biomarkers in traumatic brain injury. We investigated this possibility with a novel data set collected from neurosurgical trauma patients who presented to a level one trauma center in an urban setting. As our standard of practice, patients presenting with TBI often undergo intracranial electrode placement for the purposes of improved seizure monitoring (Marcellino et al. 2018; Waziri et al. 2009; Won et al. 2023). Patients with head injuries who presented with an initial Glasgow Coma Scale (GCS) of 8 or less received quad-lumen intracranial bolts for multimodality monitoring including depth electrodes. Likewise, patients who presented with acute subdural hemorrhages secondary to head trauma were taken to the operating room for evacuation, and subdural strip electrodes were placed intraoperatively. In either case, we were afforded the opportunity to record intracranial EEG (iEEG) signals in the acute phase in patients who sustained traumatic brain injuries.

By analyzing spectral features of these iEEG signals, we found that GCS was significantly correlated with the amount of beta power across all patients in a heterogeneous TBI population. Improvement of GCS during recovery coincided with the recovery of beta power over time, whereas beta power was usually persistently low in patients with poor neurological outcomes. Further, by analyzing how beta oscillations were distributed over space, we found that they manifest as traveling waves across the cortical surface. Finally we show that survival among TBI patients was significantly related to the amount of beta power and traveling waves observed during the recording period. Taken together, we assert that invasive recordings are a powerful tool for neuroprognostication in TBI via the presence and recovery of beta oscillatory phenomena.

## Results

We examined iEEG in 16 patients (4 female; 50.9 ± 4.4 years; mean ± SEM) who presented with traumatic brain injury. Depending on the clinical scenario, patients either underwent multimodality monitoring with a quad-lumen intracranial bolt with depth electrode (N = 7) or craniotomy with subdural strip electrode placement (N = 9) (Fig. 1a). We extracted spectral features using wavelet convolution in order to measure time varying oscillatory power in different frequency bands (Vaz et al. 2019; Addison 2017) (Fig. 1b). Changes in narrowband oscillatory power can be obscured by broadband shifts in power which are unrelated to the features of interest (Vaz et al. 2017; Cole and Voytek 2017), and we consequently utilized a previously described methodology for parameterizing spectra into periodic and aperiodic components (Donoghue et al. 2020). We define the oscillatory power of a given frequency band as the maximum difference between the periodic fit peak and the aperiodic slope (Fig. 1c).

**Figure 1.**
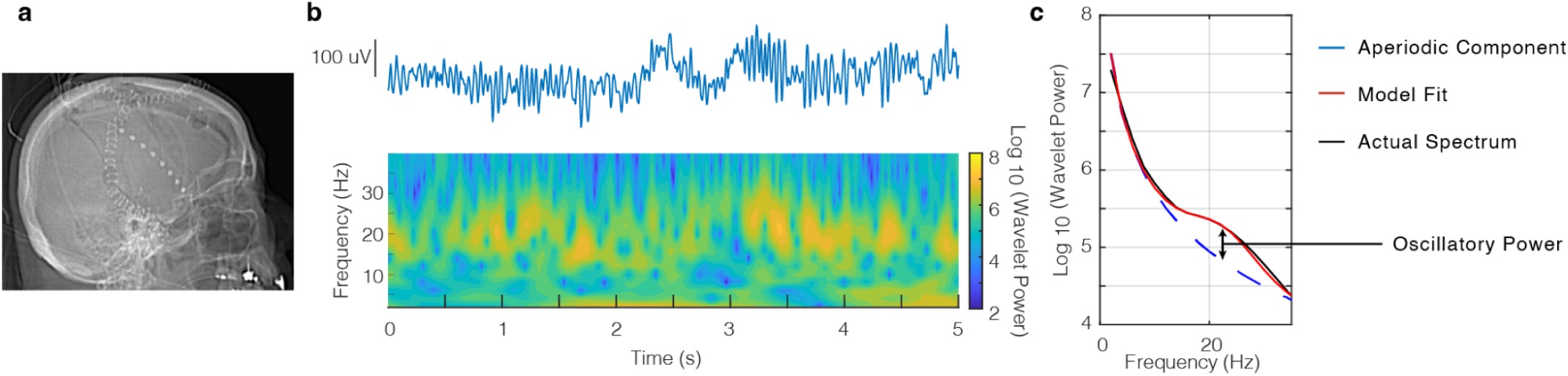
Invasive recordings in TBI patients with power spectral decomposition. a) Postoperative radiograph showing craniotomy with subdural strip electrode placement. b) Example iEEG signal obtained from an electrode (top) with wavelet spectral decomposition showing prominent beta power (bottom). c) Corresponding power spectral density (black) with full model fit (red) and aperiodic component (blue). We define beta oscillatory power as the maximum difference between the full model fit and aperiodic component.

Given preliminary scalp EEG evidence for decreased beta power (12-30 Hz) in stroke and traumatic brain injury (Wu et al. 2016; Mofakham et al. 2022), we measured these spectral features using our invasive monitors in the post-injury period. Neurological status was measured using the standard Glasgow Coma Scale (GCS) and documented on a daily basis. A representative patient who presented with head trauma and acute subdural hemorrhage (Fig. 2a) had a gradual improvement in neurological status over time where her initial GCS of 10 improved to 15 by postoperative day 3. Power spectra measured by the intraoperatively placed subdural strip electrode demonstrated a concomitant increase in oscillatory beta power corresponding to this recovery of neurological exam (Fig. 2b). Conversely, a patient who presented with multicompartmental hemorrhage including large left frontal contusion did not recover beta oscillatory power over his hospital course, and ultimately passed after care was withdrawn due to poor neurological prognosis (Fig. 2c, d).

**Figure 2.**
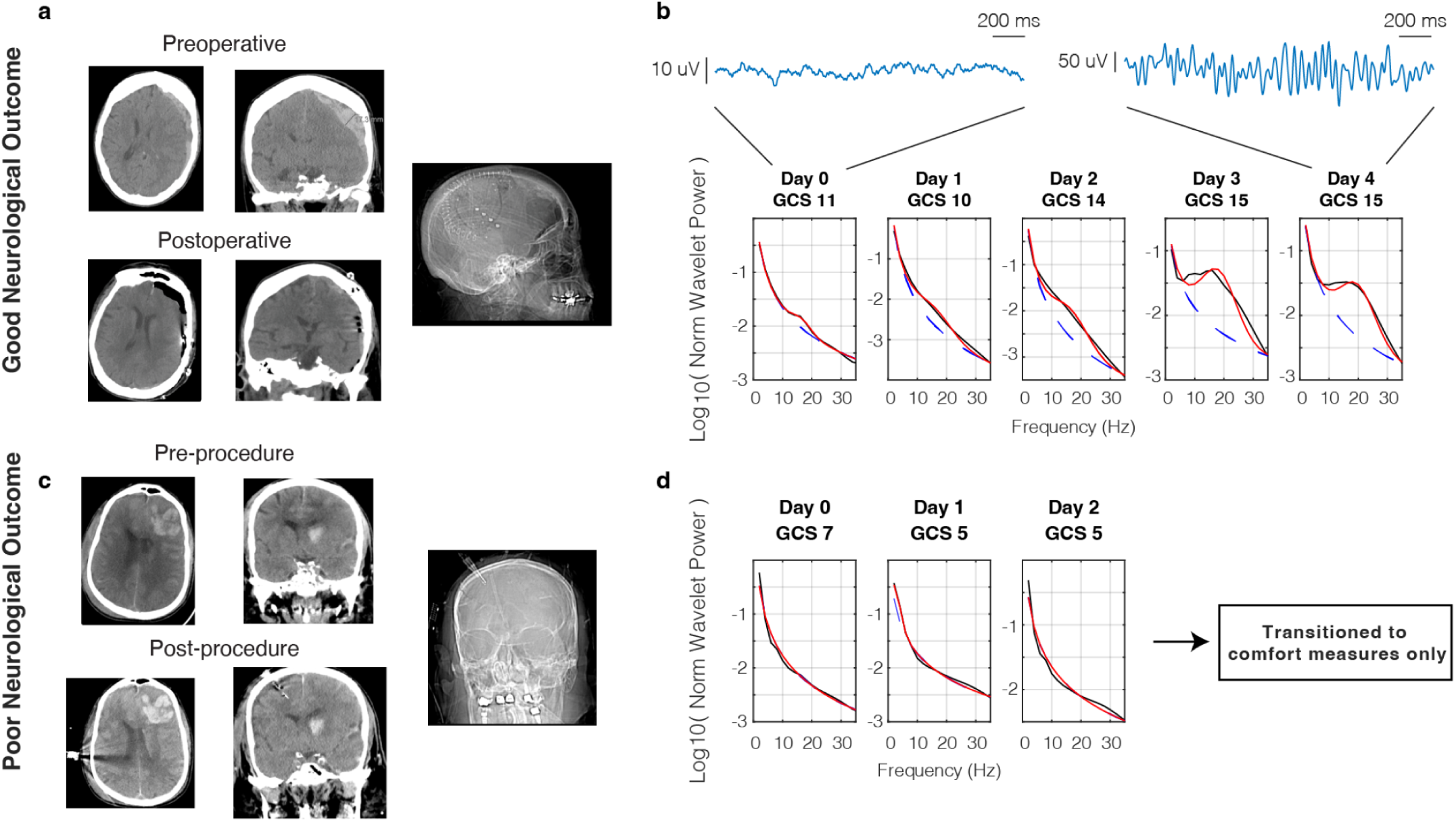
Return of beta oscillatory power with recovery from traumatic brain injury. a) Pre- and postoperative CT images of a patient with good neurological outcome who presented with acute traumatic subdural hemorrhage and was implanted with a subdural strip electrode. b) Corresponding normalized power spectral densities over time showing return of beta oscillatory power with recovery of GCS. Example raw traces are shown from an electrode that showed a significant increase in beta oscillations from day 0 to day 4. c) Pre- and postoperative CT images of a patient with poor neurological outcome who presented with blunt TBI and underwent quad-lumen intracranial bolt placement with depth electrode. d) Corresponding power spectral densities over time showing persistent deficiency of beta oscillatory power.

To quantify the relationship between beta power and GCS within individual patients with different numbers of recording sessions (3.50 ± 0.50 sessions per patient), we employed a linear mixed effects model to predict beta power with GCS and patient identity as random and fixed variables respectively (see Methods). GCS was a significant predictor of beta power across all patients (t(54) = 3.15, p = 0.003). We also compared the average GCS across days for each patient against the average intracranial beta power for all patients and found a significant correlation (r = 0.603, p = 0.013; Fig 3a). We additionally replicated this analysis using only the first or last day of the data instead of averaging across all sessions (Fig. S1). Because this data was derived from two separate device types, we also replicated these calculations in two subgroups (Fig. S2). Finally, we showed that the relationship between oscillatory power and GCS was selective for the beta band, as we were not able to replicate these results with either theta (4-8 Hz) or alpha (8-12 Hz) frequency bands (Fig. S3).

**Figure 3.**
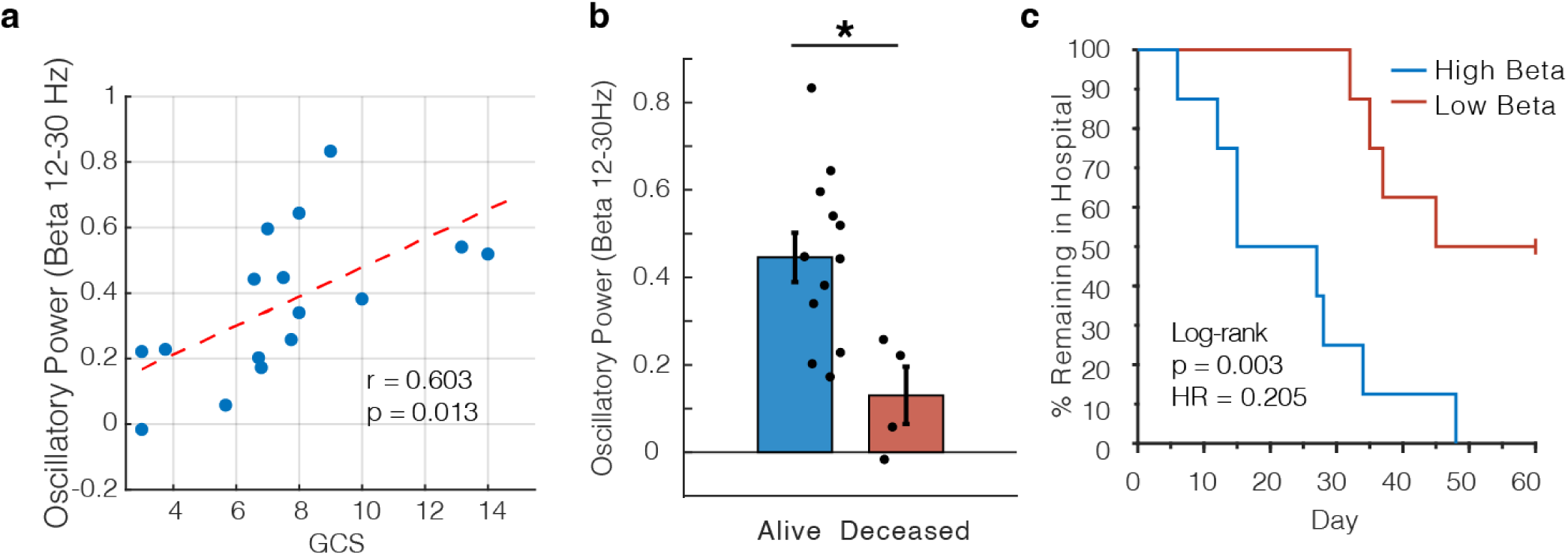
Intracranial beta oscillatory power correlates with GCS and predicts mortality and length of in-hospital stay. a) Scatter plot showing correlation between beta oscillatory power and GCS (r = 0.603, p = 0.013). Each dot represents the average values for one patient, and the red dashed line shows the line of best fit. b) Comparison of average beta oscillatory power for patients who survived their TBI versus those who died within 2 months of injury (t(14) = 2.97, p = 0.010). c) Kaplan-Meier curve showing % of patients remaining in the hospital where patients were split into two groups based on the median beta oscillatory power (log-rank test, p = 0.003).

We additionally performed analyses to rule out sedation and anesthetics in the immediate post-procedure period as a potential confounding effect of the above results. Per our clinical protocols, intubated or agitated patients typically received light sedation with propofol, fentanyl, dexmedetomidine, or midazolam. For all patients who had separate recording sessions with and without post-procedure sedation, we compared average beta power between these subgroups and found no significant difference (paired t-test between sessions within the same patients who had separate sessions with and without sedation, t(3) = 1.88, p > 0.05; unpaired t-test comparing all sessions across patients, t(18) = 1.35, p > 0.05). Moreover, we observed significant increases in post-procedure beta power in several representative patients who never received sedation (eg. Fig. 2b). Finally, we performed a separate linear mixed effects model with an interaction term between sedation and GCS to directly assess if there was a confounding effect of sedation on the amount of measured beta power, and we found that GCS was still the only significant effect (see Methods; t(52) = 2.33, p = 0.024). We assert then that the beta power effects that we report here are much more likely to be due to underlying neural dynamics than relatively small doses of sedation given in the post-procedure period.

Further, we were interested if beta oscillatory power could serve as a prognostic marker of longer term outcomes for TBI patients. We used a simple median split to divide the data into patients with high and low beta power respectively. All patient deaths in this cohort were due to withdrawal of care due to poor neurological prognosis, and we found 0% mortality in the high beta group and 50% mortality in the low beta group. We then explicitly compared the average beta oscillatory power between patients who died and those who did not, and we found that increased beta power was indeed related to survival (Fig. 3b; t(14) = 2.97, p = 0.010). We then extended this analysis with the hypothesis that high beta power would prognosticate shorter length of stay in the hospital in addition to increased mortality. We constructed Kaplan-Meier style curves for each group and found that length of stay was significantly shorter in patients with high beta power (log rank test, p = 0.003), confirming faster functional recovery times for these patients as well (Fig. 3c). We did not classify deceased patients as discharged in order to maintain a fair comparison between groups. Finally, given recent evidence suggesting that invasive recordings may offer superior detection of clinically relevant electrophysiological phenomena (Marcellino et al. 2018; Robinson, Hartings, and Foreman 2021), we directly compared these results to those obtained from analyzing simultaneously recorded scalp EEG (Fig. S4). We ran identical analyses but substituted the 8 frontal electrodes from the clinical bipolar montage. We were not able to show significance for a relationship between beta power and GCS, mortality, or length of hospital stay, suggesting that invasive measurements offer clinical advantage over traditional extracranial electrodes.

Because the beta oscillations we measure are distributed in space, we were motivated to understand if these oscillations could subserve higher order electrophysiological phenomena such as traveling waves. We leveraged a rich literature precedent showing the cognitive and sensorimotor relevance of traveling waves in the beta frequency in both humans (Takahashi et al. 2011; Stolk et al. 2019; Das et al. 2022) and nonhuman primates (Bhattacharya, Brincat, et al. 2022; Zabeh et al. 2023). Further, loss of consciousness has been linked to decreased beta traveling waves during propofol anesthesia (Bhattacharya, Donoghue, et al. 2022). Therefore we hypothesized that the beta oscillations we observed could manifest as traveling waves across the cortical surface. We adapted established procedures for extracting waves from oscillatory activity distributed across space (see Methods) (Rubino, Robbins, and Hatsopoulos 2006; Zhang et al. 2018) and found several representative examples of cortical traveling waves in patients with subdural strip electrodes (Fig. 4a). Wave rate was closely correlated to the GCS of the patients over time (Fig. 4b, top; r = 0.783, p = 0.013). We reproduced this result with a similar linear mixed effects model as above (see Methods), and GCS was a significant predictor of wave rate across patients (t(33) = 2.94, p = 0.006). Intuitively, and as previously shown (Denker et al. 2018), the rate of cortical traveling waves increased with increasing beta power (Fig. 4b, bottom; r = 0.819, p = 0.007). As with the magnitude of beta oscillatory power, we split the patient cohort by the median wave rate and found that no patients in the high wave rate group died within a 2 month period, compared to 50% of the low wave rate group. Consequently the wave rate in patients who survived their TBI was significantly higher than those who did not (Fig. 4c; t(7) = 4.91, p = 0.002). Finally, we calculated the Kaplan-Meier style curves stratified by wave rate and found again that wave rate was a predictor of discharge from the hospital, indicating that higher wave rate informs functional recovery from TBI (Fig. 4d; log rank test, p= 0.005).

**Figure 4.**
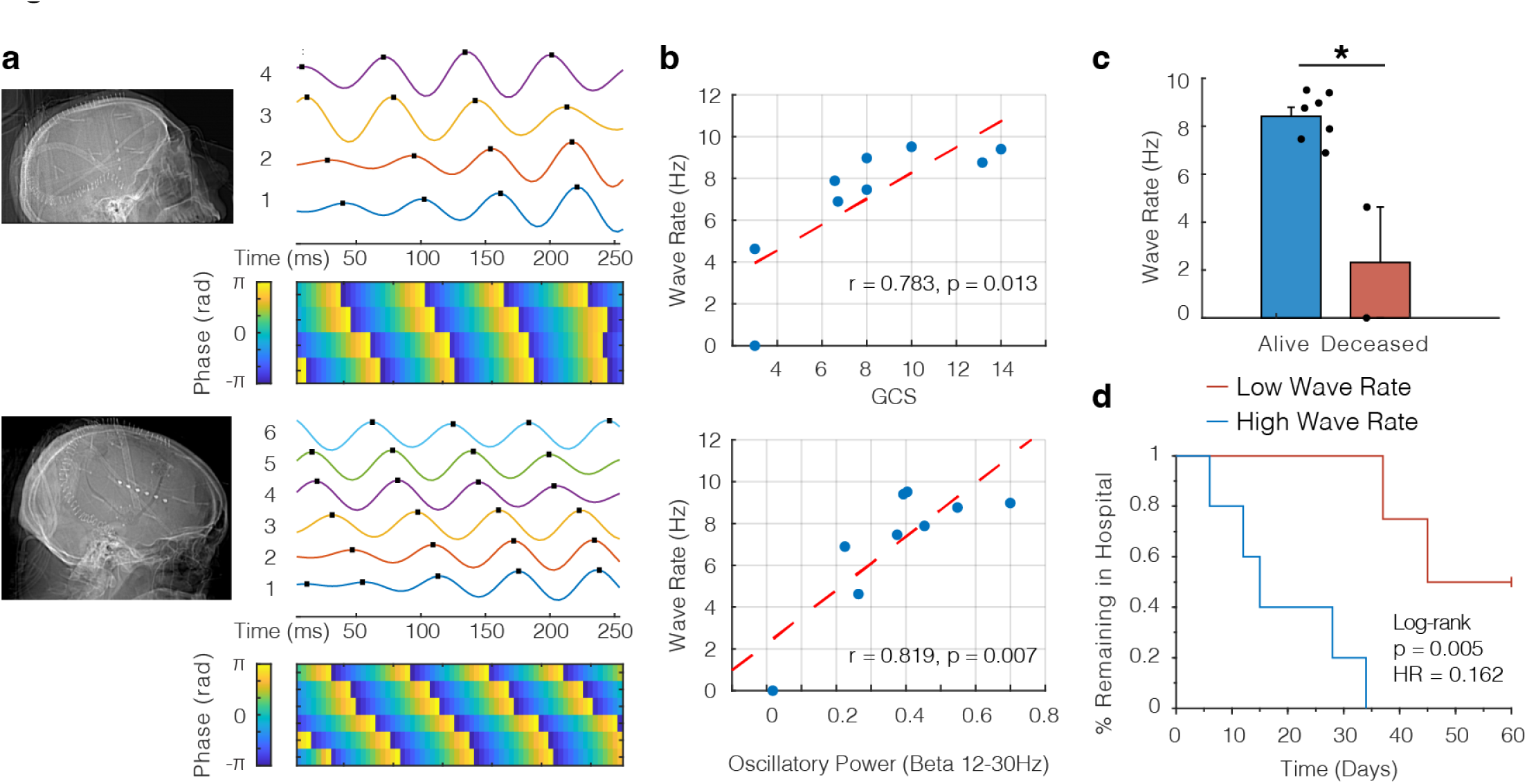
Beta oscillations manifest as traveling waves across the cortical surface. a) Two postoperative craniotomy patients who presented with acute traumatic subdural hemorrhages and were implanted with subdural strip electrodes (4 and 6 contacts respectively). In either case, robust examples of cortical traveling waves were observed. Top panels show filtered iEEG traces in the beta range (peak frequency of 16Hz; traces are shown filtered 12-20 Hz) and bottom panels show the corresponding instantaneous phase for each electrode. b) Scatter plots showing correlation between GCS and wave rate (top; r = 0.783, p = 0.013) and oscillatory power and wave rate (bottom; r = 0.819, p = 0.007). Each dot represents the average values for one patient. c) Comparison of wave rate for patients who survived their TBI versus those who died within a 2 month period (t = 4.91, p = 0.002). d) Kaplan-Meier curve showing % of patients remaining in the hospital where patients were split into two groups based on the median wave rate (log-rank test, p = 0.005). We included the median data point in the low wave rate group (given total number of patients = 9) for the purposes of this plot.

We consequently performed several control analyses to confirm that our traveling wave detection was meaningful in our data set. We first generated a null distribution of phases by randomizing values across time, and we found that the true wave rates were significantly higher than would be expected by chance (Fig. S5; t(8) = 6.74, p < 0.001). Further, we replicated our main analyses with more conservative thresholds for the duration and magnitude of phase gradient needed to classify an event as a wave (Fig. S6). As a natural control arising from this data set, we would not expect waves to travel along the axis of depth electrodes within the brain parenchyma. We therefore calculated the relationship between traveling waves along the depth electrodes and GCS and found that it was not robust across patients (Fig. S7; r = 0.50, p = 0.253). Finally, we were interested if a metric of phase gradient consistency could be used to detect waves in a similar fashion, and we were able to reproduce the relationship between wave rate and GCS and beta oscillatory power with this separately defined metric (Fig. S8).

## Discussion

Here we provide several insights into the evolution of narrowband oscillatory activity in a novel data set of invasive recordings in patients presenting with severe traumatic brain injury. We show the magnitude of beta power and the evolution of cortical traveling waves are significantly related to the neurological status of patients and that this relationship is dynamic over the course of inpatient admission. We also demonstrate that these oscillatory phenomena are tractable biomarkers of the long-term outcomes of TBI patients in that they inform mortality and ultimate hospital length of stay. Taken together, we propose that invasive recording of beta oscillations and traveling waves are efficacious biometrics for the recovery from TBI.

While we focused on the beta frequency band due to prior evidence from stroke and TBI using scalp EEG (Wu et al. 2016; Mofakham et al. 2022), we also note that the function and value of beta oscillations is well established from a neurocognitive standpoint. Beta oscillations have been heavily implicated in executive control, working memory, and sensorimotor control, particularly in the prefrontal cortex (Schmidt et al. 2019; Sherman et al. 2016). We note that our intracranial bolt depth electrode recordings were exclusively placed through burr holes above the prefrontal cortex, and subdural strips were placed along the frontoparietal convexity or superior temporal lobe, both regions previously shown to have memory related beta oscillations (Proskovec et al. 2018). It follows then that beta oscillations are a suitable biomarker for the recovery of volitional cognitive domains associated with the recovery of consciousness following TBI.

We believe that the organization of beta oscillations into cortical traveling waves is representative of the richness of spatiotemporal organization necessary for higher cognition and awareness associated with the recovery from coma. Indeed, there is an extensive literature connecting cognition, arousal, and conscious perception to beta cortical traveling waves (Takahashi et al. 2011; Stolk et al. 2019; Das et al. 2022; Bhattacharya, Brincat, et al. 2022; Zabeh et al. 2023). This line of research aligns with prior studies showing propofol induced loss of consciousness is associated with decreased beta traveling waves (Bhattacharya, Donoghue, et al. 2022) and the fragmentation of neural connectivity (Lewis et al. 2012). Together this creates a compelling argument for cortical traveling waves as a biomarker for the dynamic integration of multiple brain areas needed to sustain consciousness. Indeed this view is consistent with many reports showing decreased distributed network connectivity in disorders of consciousness (Vanhaudenhuyse et al. 2010; Demertzi, Soddu, and Laureys 2013).

Following this logic, we hypothesize that injury to thalamocortical projections is a focal point in the loss of network level activity and beta oscillatory phenomena in our cohort of TBI patients. Previous studies have proposed that the disruption of normal cortical dynamics in TBI may be due to injury to thalamo-cortical connections (Mofakham et al. 2021, 2022), and many prevailing theories of cortical wave generation rely on inputs from thalamo-cortical projections (Muller et al. 2018). Cortex is independently capable of generating intrinsic beta rhythms, but subcortical circuits involving the thalamus and basal ganglia are likely important in generating behaviorally and ecologically relevant beta oscillations (Sherman et al. 2016; Rosanova et al. 2009). Indeed, several efforts towards deep brain stimulation of thalamic nuclei for the recovery of consciousness have been undertaken with promise in rodents (Bastos et al. 2021), nonhuman primates (Tasserie et al. 2022), and humans (Giacino et al. 2012; Gummadavelli et al. 2015; Chudy et al. 2023; Schiff et al. 2023).

In summary, as our standard of practice, TBI patients undergoing either multi-modality monitoring with intracranial bolt placement or craniotomy for subdural hematoma evacuation respectively receive depth or subdural strip electrodes for improved seizure localization. We first assert that these results add to the growing volume of literature that shows improved identification of electrophysiological biomarkers with invasive recordings as opposed to scalp EEG (Marcellino et al. 2018; Robinson, Hartings, and Foreman 2021). Moving forward, we anticipate a expanded utility for invasive recordings where advances have been mostly limited to scalp EEG including cardiac arrest (Sethi et al. 2016), subarachnoid hemorrhage (Claassen, Mayer, and Hirsch 2005), and seizure monitoring after subdural hematoma evacuation (Rabinstein et al. 2010). We secondarily propose that while these electrodes are in place, features such as beta oscillatory power and traveling waves could be used to reduce prognostic uncertainty, thereby providing crucial information in end-of-life decision making (Goostrey and Muehlschlegel 2022; Fischer et al. 2022). From a practical perspective, electrophysiological biomarkers such as these could help define specific time periods for observation and provide objective metrics of clinical improvement (Miranda et al. 2023).

## Acknowledgments

We thank Casey Halpern and Iahn Cajigas for helpful and insightful comments on the manuscript. We are indebted to all patients who contributed to this study.

## Contributions

AV, CW, and DP conceptualized the study; AV performed all data analysis, software development, and visualization; KB and LB provided additional analysis code; AV, CW, SM, ST, JA, KB, SS, SA, JG, DS, JS, AR, HC, and DP curated the data; AV, CW, and DP wrote the original draft; DP supervised the study; all authors reviewed and edited the final manuscript.

## Methods

### Patients and Surgical Procedures

Patients who underwent iEEG monitoring during a three-year period between 2020 and 2022 were retrospectively identified via a prospectively maintained database of patients with cranial trauma presenting to a single level 1 trauma center located in an urban setting (Penn Presbyterian Medical Center, Philadelphia, PA). Our data set consisted of 16 patients (4 female; 50.9 ± 4.4 years) who presented with traumatic brain injury (University of Pennsylvania IRB protocol 855266).

iEEG was performed as standard of care in our center for patients with severe TBI. In patients who presented with acute subdural hemorrhage requiring craniotomy or craniectomy, a subdural strip electrode (TS08R-SP10X-000, Ad-Tech, Oak Creek, WI) was placed prior to closure. The distal lead was tunneled posterior to the incision and secured in place with suture at the skin. Per Brain Trauma Foundation guidelines, potentially salvageable patients with severe TBI (GCS < 9) underwent placement of an ICP monitor. In these patients a quad-lumen bolt (H0000-3644, Hemedex, Waltham, MA) was placed. This facilitates the placement of an ICP monitor, brain tissue oxygenation monitor, a microdialysis catheter, in addition to a depth electrode. Bolts were placed in standard fashion at or near Kocher’s point with the laterality dictated by the patient’s pathology. In these patients a 8-contact depth electrode (SD06R-SP05X-000, Ad-Tech, Oak Creek, WI) was placed through the bolt to a depth of 6cm such that the most proximal contact sat within the cortex subjacent to the bolt. In patients who received both craniotomy and bolt placement, placement of either a subdural strip or depth electrode was left to the discretion of the treating surgeon. All patients subsequently underwent postoperative non-contrast head CT.

### Intracranial EEG (iEEG) recordings

We collected intracranial EEG (iEEG) data from subdural and depth recording contacts. Subdural contacts were arranged in strip configurations with a contact radius of 1.5 mm and inter-contact spacing of 10 mm and consisted of either 4, 6, or 8 recording contacts. Depth electrodes were placed through one of the lumens of a quad-lumen intracranial bolt used for intracranial pressure monitoring, and these electrodes always had 8 contacts. At the discretion of the clinical team, iEEG signals were sampled at 256 Hz. All recorded traces were filtered with a fourth order low pass filter with cutoff frequency of 40 Hz to eliminate any possible sources of electrical line noise. For any power analyses, we re-referenced these raw signals using bipolar referencing in order to mitigate any effects of volume conduction or any biases introduced by the system hardware reference (Vaz et al. 2019). Likewise, for traveling wave analyses, we necessarily utilized monopolar referencing in order to not induce phase offsets between adjacent electrodes.

In addition to system level line noise, eye-blink artifacts, sharp transients, and inter-ictal discharges (IEDs) can generate spurious broadband power and potentially disrupt power estimates. We therefore implemented a previously reported automated event-level artifact rejection (Vaz et al. 2019). We calculated a z-score for every iEEG time point based on the gradient (first derivative), and any time point that exceeded a z-score of 5 was marked as artifactual. 100 ms before and after each identified time point was also classified as an artifact.

### Power analyses

We quantified spectral power and phase by convolving iEEG signals with complex valued Morlet wavelets (wavelet number 6) (Vaz et al. 2019; Addison 2017). We extracted one hour of data from every recording session for our analyses and discarded the first 10 minutes in order to remove line noise associated with initiation of signal recording. In all cases, we calculated spectral power using 20 linearly spaced wavelets between 2 and 40 Hz. We then squared and log-transformed the continuous-time wavelet transform to generate a continuous measure of instantaneous power. To account for different power profiles across electrodes when averaging within the same patient, we normalized the area under the power spectrum within the range of 2-40 Hz (Mofakham et al. 2022).

Narrowband spectral power changes can however be difficult to interpret, as changes in aperiodic spectra can confound these calculations (Vaz et al. 2017; Cole and Voytek 2017; Seymour, Alexander, and Maguire 2022). We therefore implemented a previously described methodology for the parameterization of aperiodic and periodic spectra in order to more accurately measure genuine beta oscillatory activity (Donoghue et al. 2020). Briefly, we modeled each power spectral density (PSD) as a sum of a general aperiodic component and multiple possible narrow-band components representing true oscillatory activity. The magnitude of difference between the aperiodic and periodic fits then gives an intuitive measure of the amount of oscillatory power that exceeds that which would be measured from a random-walk process (eg. Fig. 1c). These considerations help to safeguard against spurious conclusions about beta power that may arise from non-oscillatory phenomena (Donoghue, Schaworonkow, and Voytek 2022).

### Wave detection

We adapted established procedures for identifying traveling waves in two dimensions (Rubino, Robbins, and Hatsopoulos 2006; Zhang et al. 2018) for use with our linear arrays. We first extracted the instantaneous phase (φ) in the beta band (12-30 Hz) using the Hilbert transform of bandpass filtered signals (second order zero-phase distortion Butterworth filter). Electrodes for traveling wave analysis across each strip array were selected based on the presence of a true oscillatory peak in the frequency band of interest (Zhang et al. 2018), and phase gradients were then calculated from the these electrodes (∇φ). Importantly, the phase distribution was unwrapped in space by adding multiples of ± 2π when the jump between consecutive angles was less than π (Rubino, Robbins, and Hatsopoulos 2006). The mean phase gradient quantifies the degree of alignment of the respective gradients across electrodes as a function of time:

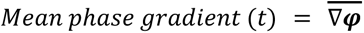

We then Z-scored phase gradient values in order to obtain normalized values for each session. Consequently, we defined a threshold of wave detection as |Z|>1 for at least 3 consecutive time points (∼12ms) in order to define the occurrence of a single wave. We note that this threshold is relatively arbitrary however, and therefore compared the number of detected waves from a randomized phase distribution to the true values and found that the empiric data always demonstrated more waves than would be expected by chance (Fig. S5). We also conducted a further control analysis where we reproduced the main results using more conservative thresholds in both duration and magnitude of phase gradient for wave detection (Fig. S6), and we showed that traveling wave effects are specific for cortical strip electrodes as opposed to parenchymal depth electrodes (Fig. S7).

Finally, we were interested if a metric of phase gradient consistency, as opposed to magnitude, could also detect traveling waves and thereby reproduce our results with a separate methodology. A consistently nonuniform distribution of phase gradients between electrodes would indicate the presence of a traveling wave. We consequently used the resultant mean vector length to quantify phase gradient consistency (equal to 0 for uniform distribution of phase gradients; (Zhang and Jacobs 2015)):

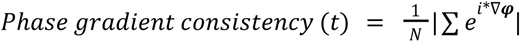

We note that this formulation is equivalent to the phase locking value (PLV; (Lachaux et al. 1999)). Using this metric, we found that traveling wave rate was again correlated with beta oscillatory power and GCS, with similar average wave rates as calculated with phase gradient magnitude (Fig. S8). Indeed the wave rates were significantly greater than would be expected by chance after shuffling phase values in time (t(8) = 4.94, p = 0.001).

We note here that the use of clinically indicated linear arrays limited our evaluation of wave directionality and therefore wave speed. Likewise our inability to detect waves traveling in a direction more perpendicular to the linear array likely resulted in a lower wave detection rate than may be in actuality. We propose then that as the indications for invasive recordings in TBI increase in the near future, grid electrode arrays may be of significantly higher clinical utility given improved detection of distributed electrophysiological activity.

### Statistical methods

To use an appropriate statistical test for the relationship over time between beta oscillatory power and GCS for data stratified by a different number of sessions for each patient, we employed a linear mixed effects model of the form:

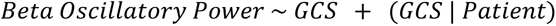

In this case, the model allows for a random intercept and slope with possible correlation between them, and the null hypothesis is that there is no relationship between GCS and beta oscillatory power. Model fits were performed with the ‘fitlme’ function in MATLAB. Likewise, we substituted wave rate for beta oscillatory power and used the same model when investigating the relationship between wave rate and GCS:

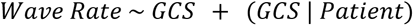

For our statistical tests concerning the significance of the difference between Kaplan-Meier curves from different patient groups, we employed the log-rank test. This test has the null hypothesis that there is no difference between populations in the probability of an event occurring over time. For example, when comparing high versus low beta oscillatory power groups, the log-rank test assessed the difference over time in discharge dates from the hospital (Fig. 3b). We calculated the log-rank test using the MatSurv function in MATLAB (Creed, Gerke, and Berglund 2020).

We further constructed a separate linear mixed effects model to account for the potential confounding effects of sedation on beta oscillatory power:

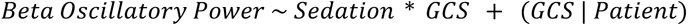

In this way, we tested explicit interaction terms related to sedation, neither of which were significant (Sedation t(52) = -0.55, p = 0.58; GCS:Sedation t(52) = -0.21, p = 0.83). Importantly, the effect of GCS was still significant in this model (GCS, t(52) = 2.33, p = 0.024), supporting the assertion that beta power in our data is most closely linked with coma status as opposed to clinically indicated sedation.

---

**Figure S1.**
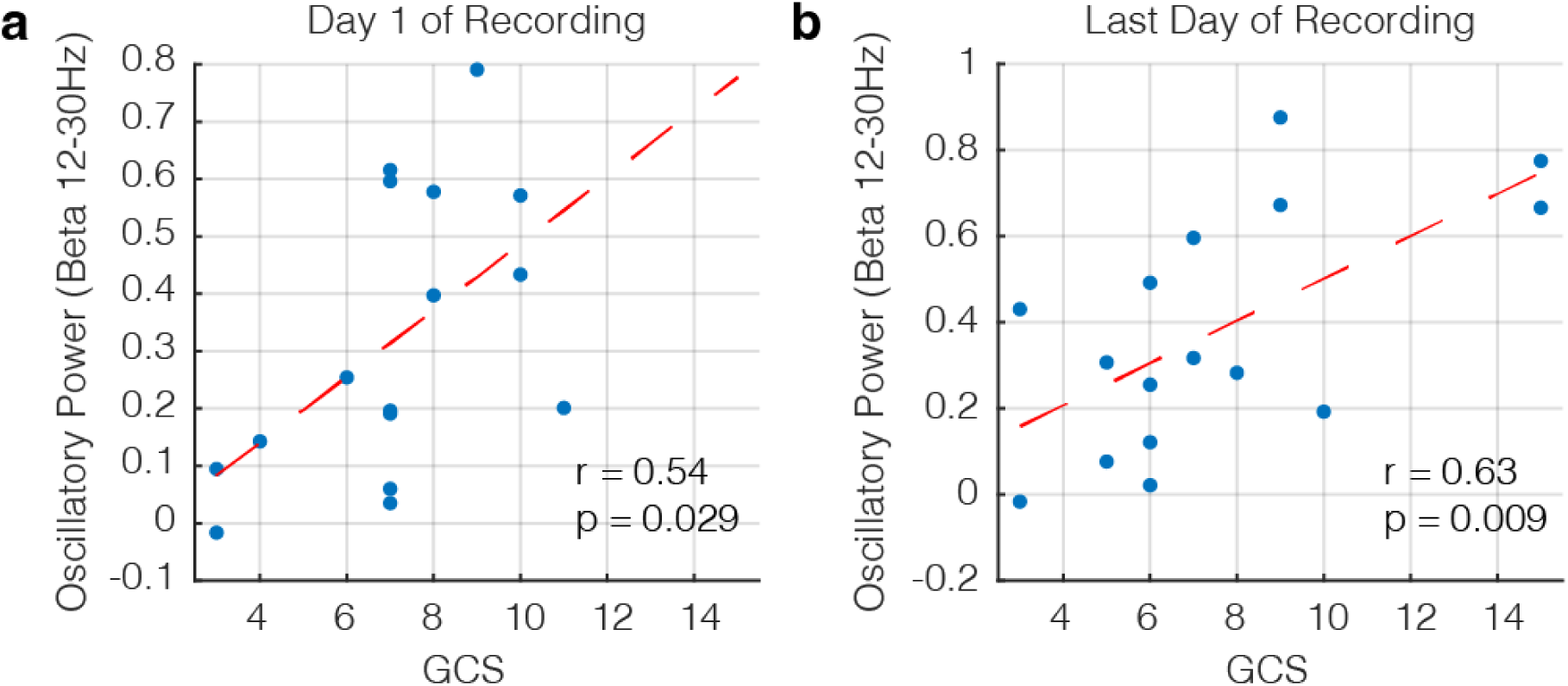
Relationship between beta oscillatory power and GCS is the same regardless of the recording day of interest. a) Scatter plot showing the correlation between beta oscillatory power and GCS for the first day of recording (r = 0.54, p = 0.029). Each dot represents the average values for one patient, and the red dashed line shows the line of best fit. b) Scatter plot showing the correlation for the last day of recording (ie. the last day before hardware was removed; r = 0.63, p = 0.009).

**Figure S2.**
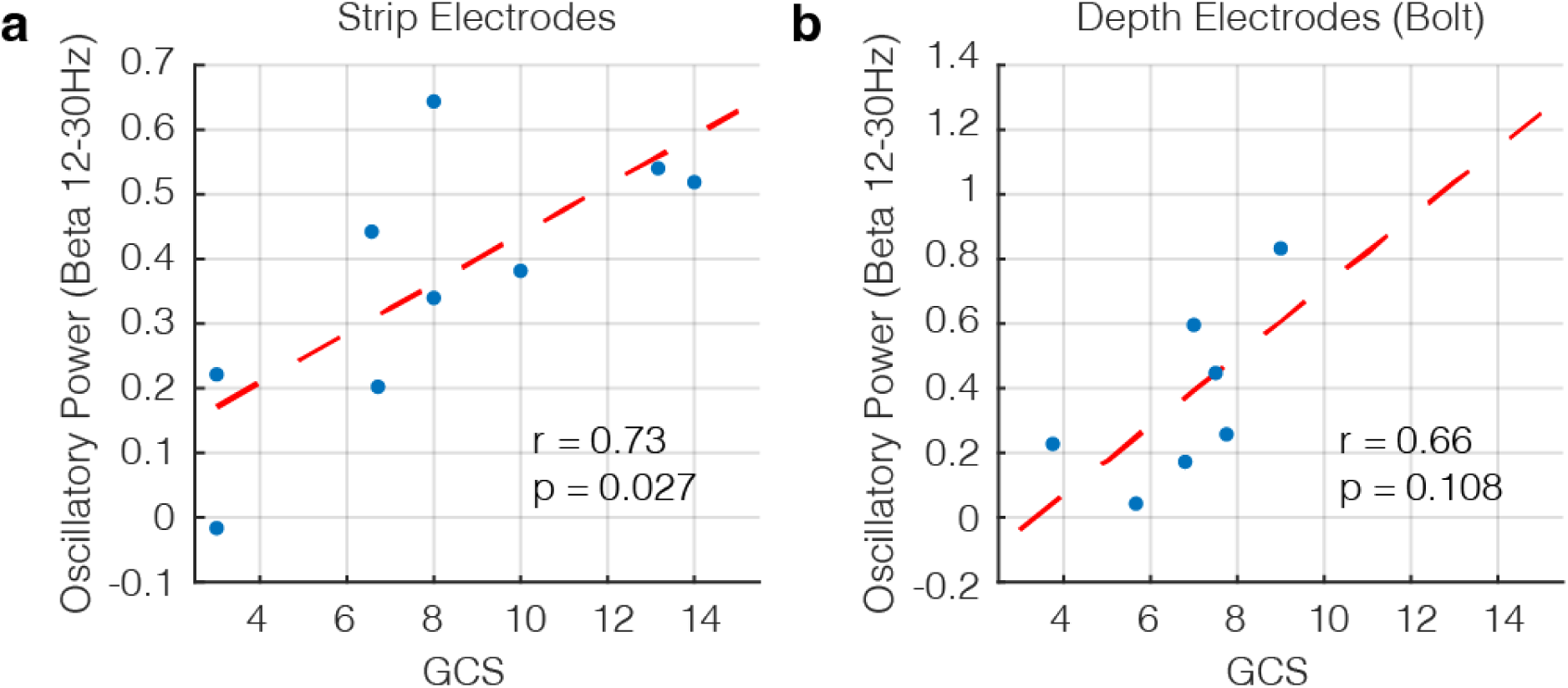
Replication of main results separately using strip or depth electrodes. a) Scatter plot showing the correlation between beta oscillatory power and GCS for patients implanted with strip electrodes after craniotomy for evacuation of acute subdural hematoma (n = 9; r = 0.73, p = 0.027). Each dot represents the average values for one patient, and the red dashed line shows the line of best fit. b) Scatter plot showing the correlation for patients implanted with depth electrodes via quad-lumen intracranial bolts after blunt TBI (n = 7; r = 0.66, p = 0.108). We note here that the overall correlation is not significant (p > 0.05), but clearly trending towards significance.

**Figure S3.**
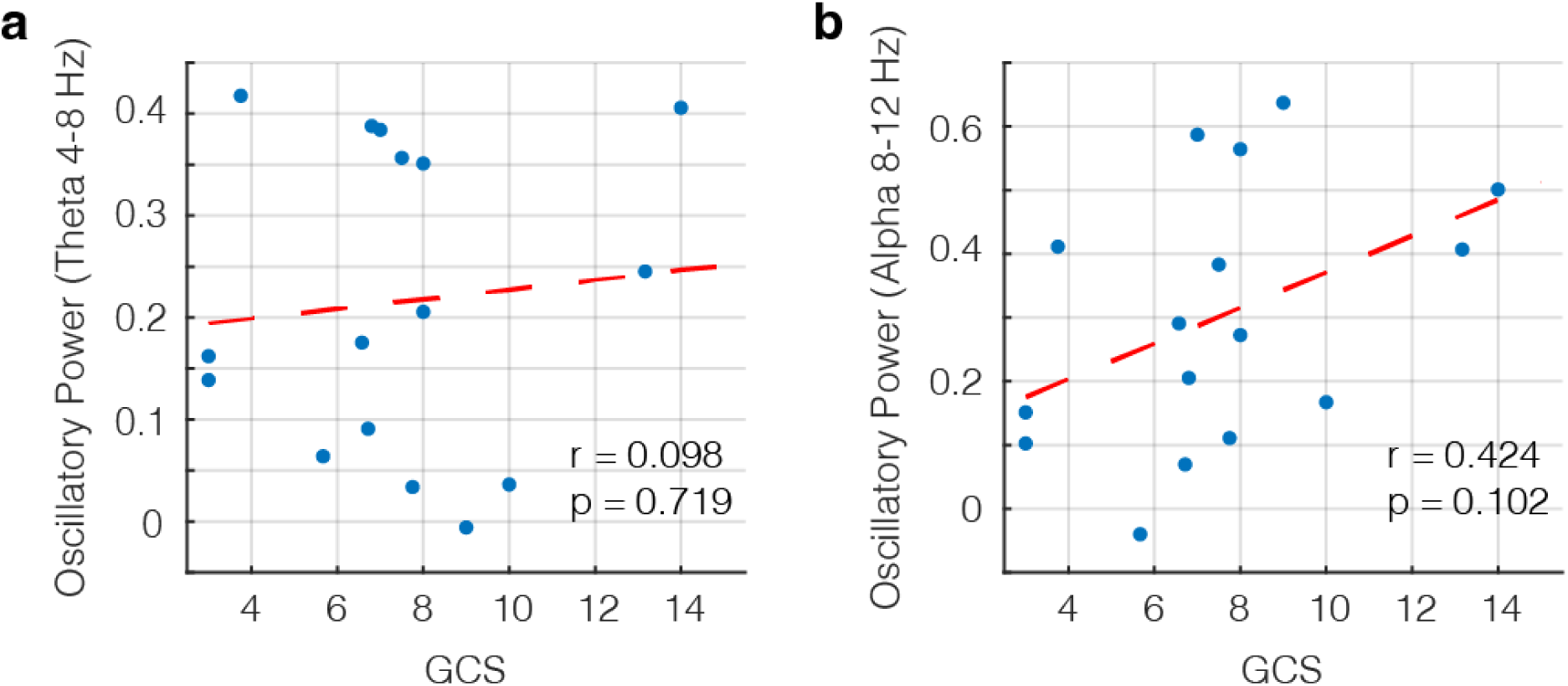
Correlation between GCS and oscillatory power is not observed in the theta or alpha frequency bands. a) Scatter plot showing the correlation between theta (4-8 Hz) oscillatory power and GCS (r = 0.098, p = 0.719). Each dot represents the average values for one patient, and the red dashed line shows the line of best fit. b) Scatter plot showing the correlation between alpha (8-12 Hz) oscillatory power and GCS (r = 0.424, p = 0.102).

**Figure S4.**
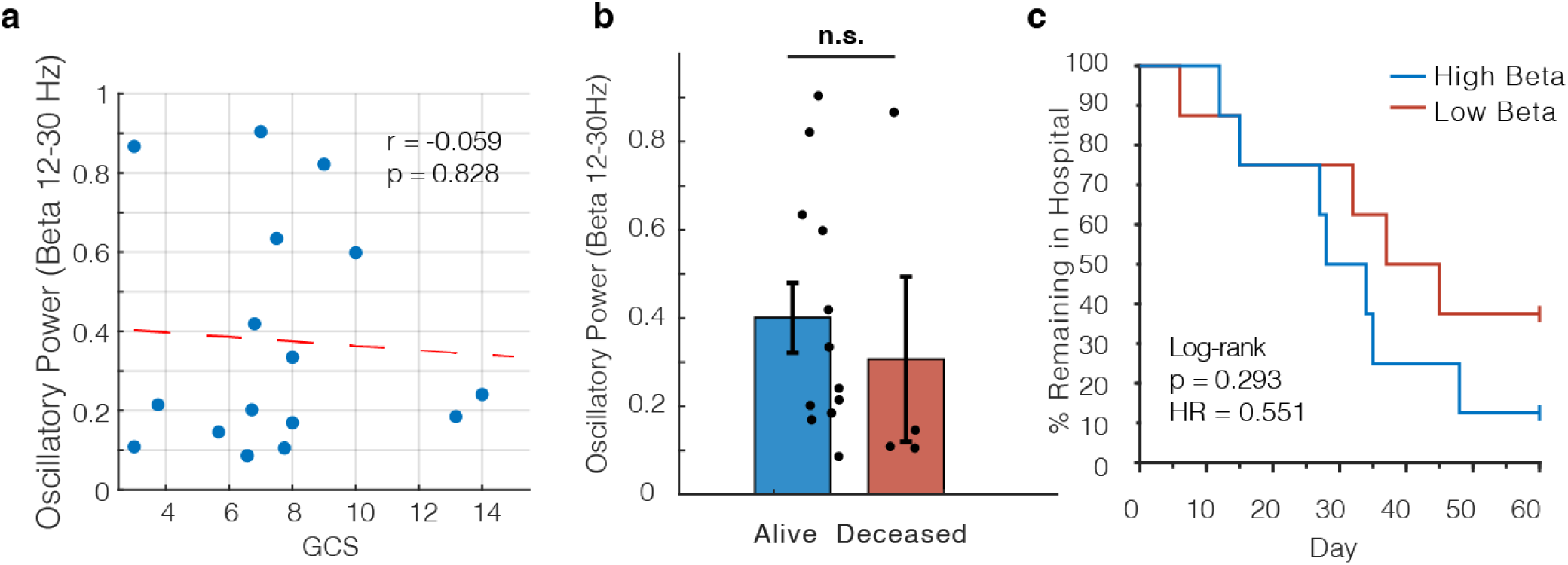
Scalp EEG beta oscillatory power does not significantly correlate with GCS or predict mortality or length of in-hospital stay. a) Scatter plot showing correlation between beta oscillatory power and GCS (r = -0.059, p = 0.828). Each dot represents the average values for one patient, and the red dashed line shows the line of best fit. b) Comparison of average beta oscillatory power for patients who survived their TBI versus those who died within 2 months of injury (p > 0.05). c) Kaplan-Meier curve showing % of patients remaining in the hospital where patients were split into two groups based on the median beta oscillatory power (log-rank test, p = 0.293).

**Figure S5.**
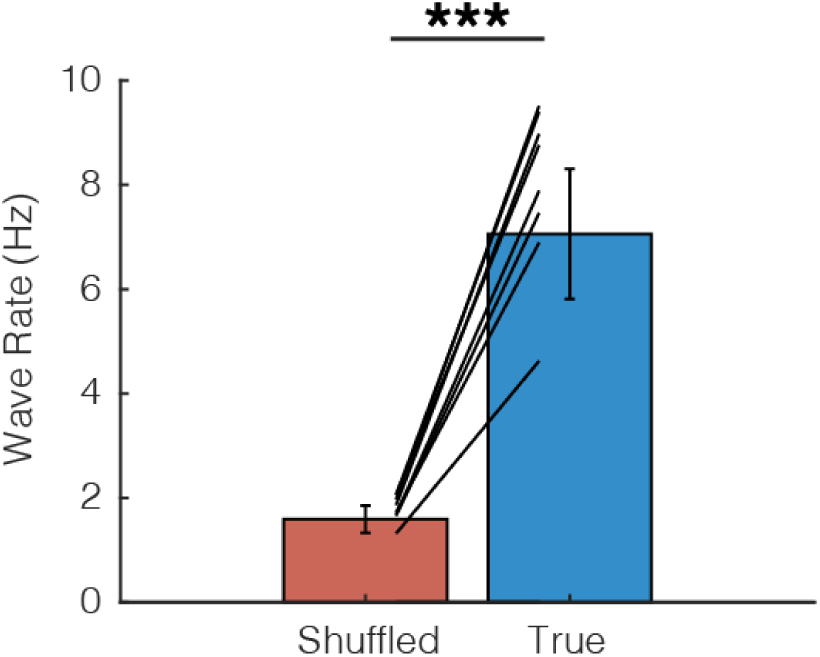
Control analysis comparing true wave detection rate versus chance. Shuffled distributions were generated by randomizing phase values in time and recalculating the wave detection as in the true case. In all patients (except in one patient where no waves were detected in either case), wave rate was higher in the true data than in the shuffled distribution, indicating that our wave detection methodology detects genuine oscillatory phenomena (t(8) = 6.74, p <0.001).

**Figure S6.**
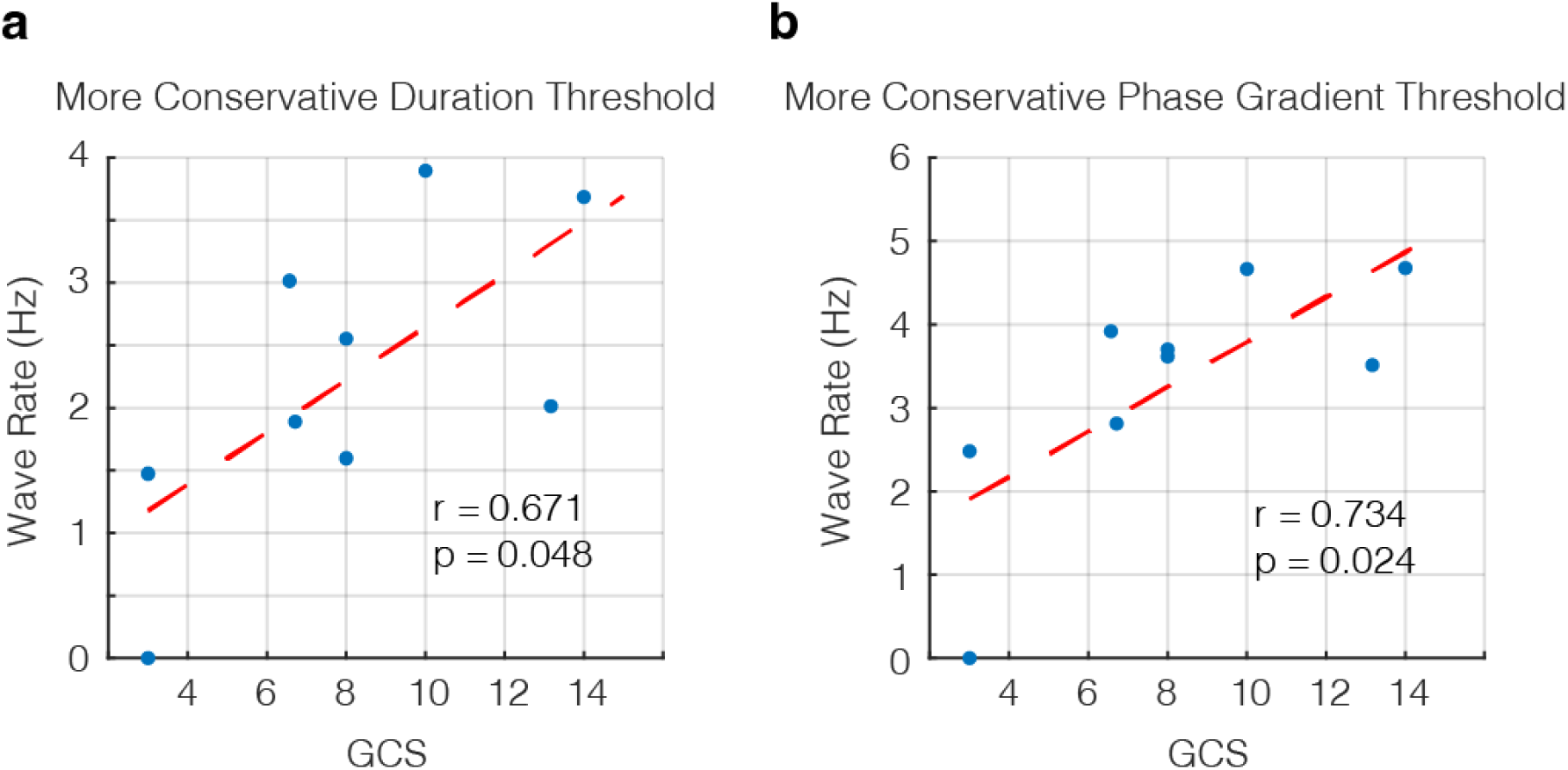
Replication of wave rate results with more conservative detection thresholds. a) Correlation between wave rate and GCS with wave detection threshold increased to 8 time points (31.25 ms, roughly equivalent to one cycle at 30Hz; r = 0.671, p = 0.048). Each dot represents the average values for one patient, and the red dashed line shows the line of best fit. b) Correlation between wave rate and GCS with wave detection threshold increased to |Z| > 1.5 (r = 0.734, p = 0.024).

**Figure S7.**
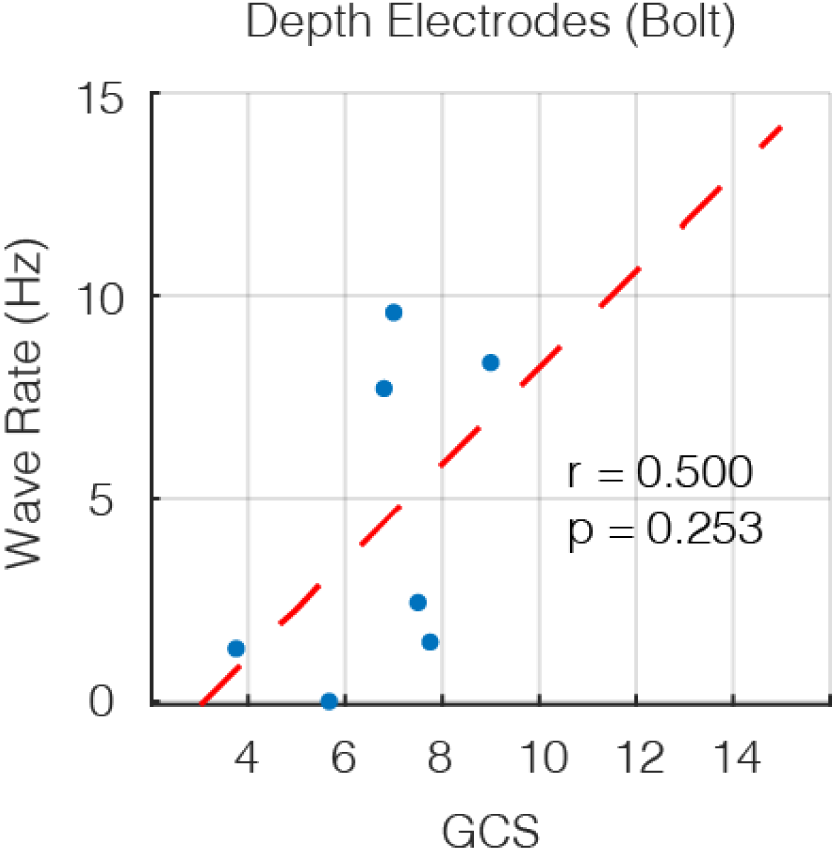
Wave rate and GCS are not significantly correlated in patients implanted with depth electrodes via quad-lumen intracranial bolts. Each dot represents the average values for one patient, and the red dashed line shows the line of best fit (r = 0.500, p = 0.253). We infer that the lack of significant correlation is due to the difficulty in detecting traveling cortical waves along the course of depth electrodes that are inserted deep within the parenchyma.

**Figure S8.**
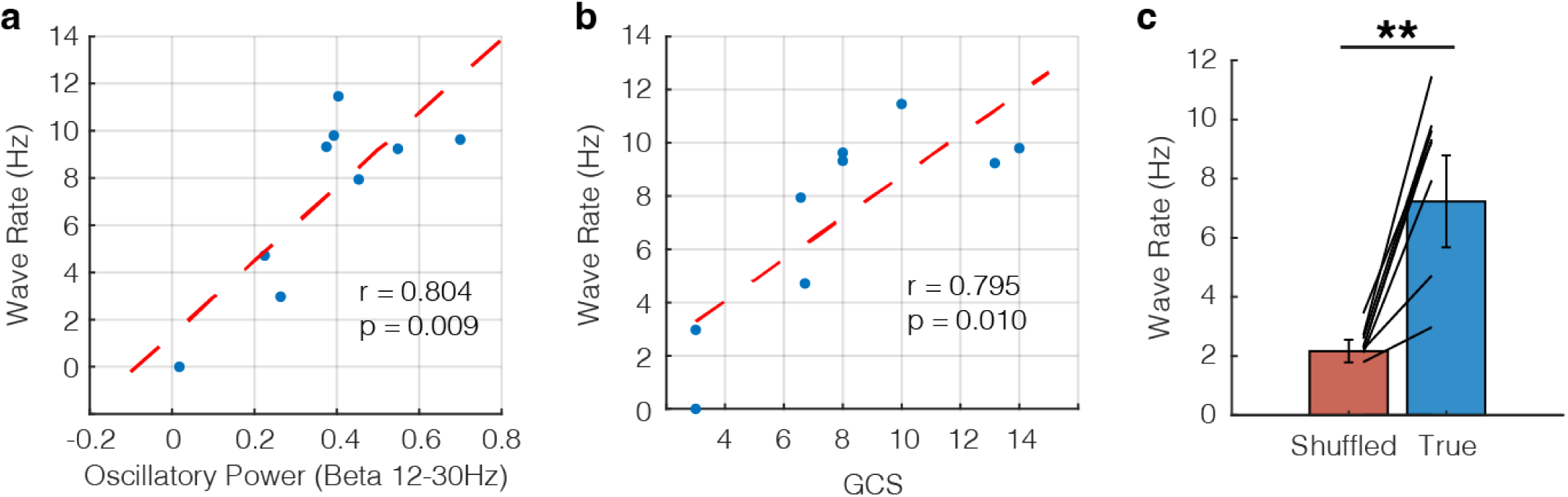
Replication of wave rate results with phase gradient consistency. a) Correlation between wave rate and beta oscillatory power as detected with phase gradient consistency (r = 0.804, p = 0.009). Each dot represents the average values for one patient, and the red dashed line shows the line of best fit. b) Correlation between wave rate and GCS (r = 0.785, p = 0.010). c) Control analysis comparing true wave detection rate versus chance (as obtained by randomizing phase values in time). In all patients (except in one patient where no waves were detected in either case), wave rate was higher in the true data than in the shuffled distribution (t(8) = 4.94, p = 0.001).

**Table S1.**

| Patient | Age | Gender | Mechanism | Diagnosis | Surgery / Device | In Hospital Mortality |
| --- | --- | --- | --- | --- | --- | --- |
| 1 | 71 | M | Found down | mICH | R frontal bolt, EVD | Yes<br>Comfort care →<br>Palliative<br>Extubation Day 6 |
| 2 | 85 | F | MVC | SDH | R craniotomy, subdural strip electrode | No<br>Discharged<br>POD15<br>GCS15 |
| 3 | 58 | M | Fall | mICH | R frontal bolt | Yes<br>Comfort care →<br>Palliative<br>extubation Day 5 |
| 4 | 30 | M | GSW | mICH | Bifrontal craniectomy, EVD, subdural strip electrode | No<br>Discharged<br>POD34<br>GCS15 |
| 5 | 69 | M | Fall | mICH | R craniotomy, Revision R craniotomy POD3 with strip electrode, bilateral MMA embolization | No<br>Discharged<br>POD12<br>GCS15 |
| 6 | 58 | M | Found down | R SDH | R craniotomy, subdural strip electrode, L frontal bolt | No<br>Discharged<br>POD28<br>GCS15 |
| 7 | 30 | M | Fall | L SDH / EDH | L craniotomy, R frontal bolt | No<br>Discharged<br>POD35<br>GCS15 |
| 8 | 68 | M | Fall | mICH | R craniotomy, subdural strip electrode | No<br>Discharged<br>POD45<br>GCS14 |
| 9 | 59 | M | Fall | L SDH | L craniotomy, subdural strip electrode | Yes<br>Comfort care →<br>Palliative<br>extubation Day 2 |
| 10 | 65 | M | MVC | mICH | L craniotomy, R frontal bolt, R frontal EVD | No<br>Discharged<br>hospital day 32<br>GCS14 |
| 11 | 33 | F | MVC | mICH | R frontal bolt | No<br>Discharged<br>hospital day 27<br>GCS15 |
| 12 | 53 | M | Fall | L SDH | L craniotomy, subdural<br>strip electrode | Yes<br>Comfort care<br>POD2 |
| 13 | 28 | M | GSW | mICH | R craniotomy, subdural<br>strip electrode, later L<br>frontal bolt | No<br>Discharged<br>POD37 |
| 14 | 33 | M | GSW | mICH | L craniotomy, R frontal<br>bolt | No<br>Discharged<br>POD48 |
| 15 | 74 | F | Fall | L SDH | L craniotomy, subdural<br>strip electrode | No<br>Discharged<br>POD6 |
| 16 | 27 | F | MVC | L SDH /<br>EDH | L skull fracture elevation,<br>R frontal bolt | No<br>Discharged<br>POD15<br>GCS15 |

SDH - Subdural hematoma
EDH - Epidural hematoma
mICH - Multicompartmental intracranial hemorrhage
MVC - Motor vehicle collision
GSW - Gunshot wound
EVD - External ventricular drain

